# Emergent feasibility in random ecological systems with higher-order interactions

**DOI:** 10.64898/2026.06.11.728491

**Authors:** Pablo Lechón-Alonso, Alexander Strang, Paul Breiding, Stefano Allesina

## Abstract

A recurring lesson from random ecological models is that coexistence is hard to come by: in the Generalized Lotka-Volterra (GLV) model with pairwise interactions, the probability that randomly sampled parameters admit a positive (feasible) equilibrium – a necessary condition for coexistence – is exactly 1*/*2^*n*^ in *n* species, vanishing rapidly with diversity. This rarity is often read as evidence that coexistence demands specific ecological mechanisms. Real interactions, however, are rarely strictly pairwise: any nonlinear dependence of one species’ growth rate on another’s abundance, Taylor-expanded, generates higher-order interactions (HOIs) of increasing degree. Treating the interaction order *d* as a knob that indexes this nonlinearity, we map the random GLV with HOIs onto the Kostlan-Shub-Smale class of random polynomial systems and approximate the probability of feasibility (*P*_*f*_ ) analytically. We find a phase transition at *d* = 4: below this threshold, *P*_*f*_ decays with diversity as in the pairwise case; above it, the exponential proliferation of equilibria outpaces the probability that any given equilibrium is feasible, and the probability of feasibility increases with *n*, approaching one. The transition appears to be universal across symmetric coefficient distributions, but vanishes when sign symmetry of the parameter distribution is broken. This work uncovers a route by which feasibility emerges from nonlinearity alone, with no fine-tuning of parameters and no appeal to specific ecological mechanisms.

## INTRODUCTION

Ecological theory has long taken the species pair as its unit of analysis, and much of our intuition about the co-existence of many species descends from pairwise models – the Generalized Lotka-Volterra (GLV) model prominent among them. Drawing the parameters at random allows to probe the typical behavior of a structureless community against which coexistence can be judged [1]. Randomly parametrized pairwise models share a common property: at high diversity they are very unlikely to admit equilibria in which all species are present at positive abundance [2–14] (feasible equilibria). Feasibility is easily destroyed by adding species, because random pairwise models lack the ecological mechanisms (self-regulation, positive intrinsic growth rates, predator-prey structure, etc.) that are thought to prevent extinctions.

Species interactions, however, often involve complexities that cannot be captured through the lens of the species pair – spatial and structural complexity [15–17], behavior switching [18–21], habitat modification [22, 23], non-linearities in prey consumption [24, 25], evolution [26]. For example, a caged predator – able to emit cues but unable to consume – can reshape the competition between two prey sharing a resource: the threat of predation alone suppresses their foraging, altering how each depletes the resource and thereby modifying the interaction between them [20]. On a smaller scale [22], the bacterium *Escherichia coli* can successfully invade cultures of *Chlamydomonas reinhardtii* as well as cultures of *Tetrahymena thermophila*, but it cannot invade a co-culture of the two, because *C. reinhardtii* inhibits *E. coli* aggregation specifically in the presence of *T. thermophila*, exposing the bacterium to predation [23]. Interactions shaped by these mechanisms are better understood through the lens of *higher-order interactions* (HOIs): the interaction between two species is modulated by a third (three-way interactions), which may itself be modulated by a fourth, and so on. Beyond such literal multi-species mechanisms, any smooth nonlinear interaction is, after Taylor expansion around an operating point – rendering HOIs a generic, agnostic representation modelling highly nonlinear systems [14, 27, 28].

Although their existence and relevance were initially debated [18, 28–33], HOIs are now an active area of research in both experimental [22–24, 34–43] and theoretical [44–49] ecology. Their analysis, however, is hard: explicit steady-state solutions of systems with HOIs do not exist in general [50], hindering the standard first step in analysis. As a result, existing feasibility results are confined to three- or four-way interactions, treated using mathematical heuristics [48], computer simulations [14, 44], or reduction to effective pairwise interactions [45].

AlAdwani and Saavedra delivered the first results on feasibility under generalized HOIs, showing numerically that the probability of feasibility increases with interaction order [46, 47, 51]. How that probability scales jointly with diversity and interaction order, however, has remained out of reach. Recent work [52] shows that a random replicator equation maps to a well-studied family of random polynomials, the Kostlan-Shub-Smale (KSS) ensemble [53, 54], reducing the counting of equilibria to the counting of polynomial zeros. Because replicator and GLV dynamics are formally equivalent [55–57], this mapping carries over directly to GLV with HOIs, putting the tools of random polynomial theory also at the disposal of ecology; in analogy to how random matrix theory has shaped our thinking about pairwise systems [58, 59].

With this machinery, we ask how does the probability of feasibility scale with diversity once interactions are allowed to be higher-order? We find that below a critical interaction order *d* = 4, the probability of feasibility decays exponentially with the number of species, recovering the familiar intuition from pairwise theory. Strikingly however, beyond *d* = 4, probability of feasibility grows with diversity and rapidly approaches one. Simulations suggest that the transition is universal accross distributions that are symmetric around 0, and breaks down when symmetry in the parameter distribution is removed. This work suggests that feasibility (a necessary condition for coexistence) can emerge without fine-tuned constraints, as a consequence of nonlinearity combined with statistical symmetry.

## HIGHER-ORDER INTERACTIONS PROMOTE FEASIBILITY

We study the per-capita generalized Lotka-Volterra dynamics in which species interact through terms of increasing order [31],

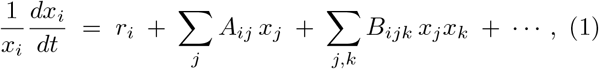

where *r*_*i*_ is the intrinsic growth rate of species *i, A*_*ij*_ the pairwise interaction coefficients, *B*_*ijk*_ the three-way coefficients, and the ellipsis collects interactions of still higher order. We let *d* denote the highest degree appearing on the right-hand side so that *d* = 1 recovers the pairwise GLV, *d* = 2 adds three-way interactions, and general *d* collects all interactions up to degree *d*. We index a given interaction by its degree *p*, the number of abundances it multiplies: *p* = 0 for the growth rate, *p* = 1 for the pairwise couplings *A*_*ij*_, *p* = 2 for the three-way terms *B*_*ijk*_ *x*_*j*_*x*_*k*_, and so on up to *p* = *d*. Equilibria are the zeros of the polynomial system in Eq. (1), *f*_*i*_(*x*) = 0; feasible equilibria are those with *x*_*i*_ *>* 0 for all *i*.

We draw the interaction coefficients randomly, each an independent mean-zero Gaussian. Setting the variance of every degree-*p* coefficient to 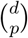 places the dynamics in the Kostlan-Shub-Smale (KSS) ensemble of random polynomial systems [53]: the per-capita rates are then equivalent to *n* homogeneous polynomials in *n* + 1 variables of common degree *d* (see Methods and SI Sec. VI for the mapping, and [52] for the original argument in the context of multiplayer evolutionary games).

A degree-*d* system of this kind has *d*^*n*^ complex roots by Bézout’s theorem; the expected number of these that are real is

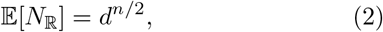

the Kostlan-Shub-Smale-Edelman identity [53]. None of these roots sits at infinite abundance, so the count carries over to the real space ℝ^*n*^ in which equilibria live (see Methods).

The real roots distribute uniformly across the 2^*n*^ orthants of ℝ^*n*^ in expectation, and because of this, we show (see Methods, and [52]) that the expected number of feasible equilibria is

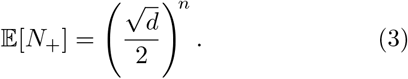

As a function of diversity *n*, E[*N*_+_] is monotonically decreasing for *d <* 4, identically equal to one for *d* = 4, and monotonically increasing for *d >* 4 (SI, Sec. V). The same counting applies when species carry different interaction orders *d*_*i*_: the expected number of feasible equilibria becomes (*d*_1_*· · · d*_*n*_)^1*/*2^*/*2^*n*^, and feasibility still emerges whenever the product *d*_1_ *· · · d*_*n*_ outgrows 4^*n*^ (SI Sec. X). The ecologically relevant quantity, however, is not the expected count of feasible equilibria but the probability *P*_*f*_ that at least one exists. Approximating root signs as independent across the 2^*n*^ orthants; we can upper bound the probability of feasibility (see Methods) as

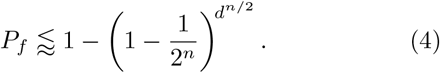

Equation (4) is a Jensen upper bound on the independence approximation, not on *P*_*f*_ itself; empirically it tracks the simulated *P*_*f*_ closely (Fig. 1**a**). At *d* = 1 this collapses to *P*_*f*_ = 1*/*2^*n*^, the pairwise baseline of Serván *et al*. [9]. For *d <* 4 the exponent *d*^*n/*2^ grows more slowly than 2^*n*^ and *P*_*f*_ vanishes exponentially in *n*; for *d >* 4 the exponent outruns 2^*n*^ and *P*_*f*_ *→* 1. The line *d* = 4 is the phase boundary, on which the two rates are matched. Numerical simulations using homotopy continuation [14, 60] agree with Eq. 4 across *n* and *d* (Fig. 1**a**); the corresponding (*n, d*) phase diagram in Fig. 1**b** makes the transition visible at a glance, with feasibility collapsing under diversity below *d* = 4 and emerging from it above. The approximation provably converges to 1 for *d >* 4; the emergence claim therefore rests on the agreement between Eq. (4) and simulations, which we probe further in the next section.

**FIG. 1.**
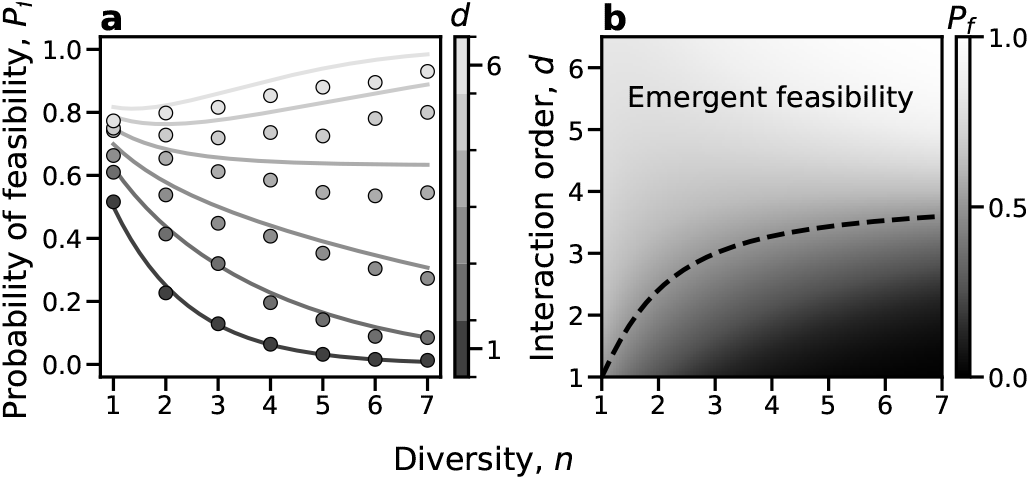
Phase transition in feasibility at *d* = 4. (**a**) Probability of feasibility *P*_*f*_ versus diversity *n*, from the independence approximation in Eq. 4 (curves, colored by interaction order *d*) and from homotopy continuation simulations of the random GLV+HOI system (points). *P*_*f*_ decays with *n* for *d <* 4 and grows with *n* for *d >* 4. (**b**) Phase diagram in the (*n, d*) plane: heatmap of *P*_*f*_ from the analytic expression. The dashed line marks the contour line *P*_*f*_ = 0.5

### UNIVERSALITY AND THE ROLE OF SIGN SYMMETRY

The KSS ensemble of the previous section served as a theoretical baseline, but as an ecological model it is unrealistic for several reasons. First, the coefficients *A*_*ij*_, *B*_*ijk*_, etc have a binomial variance scaling 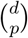 that peaks at *p* = *d/*2, meaning that intermediate-order interactions are the most variable component of the dynamics. There is no biological reason higher-order terms should be amplified relative to growth rates and pairwise couplings. Second, the KSS ensemble is dense: every interaction coefficient is non-zero. In addition, this result rests on the assumption of Gaussianity, but the question of whether it holds for a broader family of distributions remains. We relax each property in turn and ask whether the transition at *d* = 4 maintained. As we show below, the transition survives the first three relaxations, but breaks when relaxing symmetry (SI Appendix).

Reading Eq. (1) literally as a multivariate Taylor expansion of a smooth per-capita dynamics around an operating point, *f*_*i*_(*x*) = ∑_*l*_ *a*_*l*_ *x*^*l*^ with *a*_*l*_ = (*∂*^*l*^*f*_*i*_)(0)*/l*! And i.i.d. Gaussian derivatives (*∂*^*l*^*f*_*i*_)(0) yields the coefficient variance (see Methods)

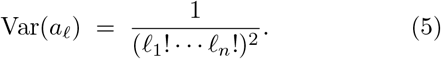

Higher-order monomials are further suppresed the higher the interaction order, encoding the intuition that nonlinearity is a perturbation of the linearized dynamics.

Numerical estimates of *P*_*f*_ under this variance fall on the same side of *d* = 4 as the KSS baseline (crosses, Fig. 2**a**): *P*_*f*_ still decays with *n* for *d <* 4, and grows with *n* for *d >* 4.

**FIG. 2.**
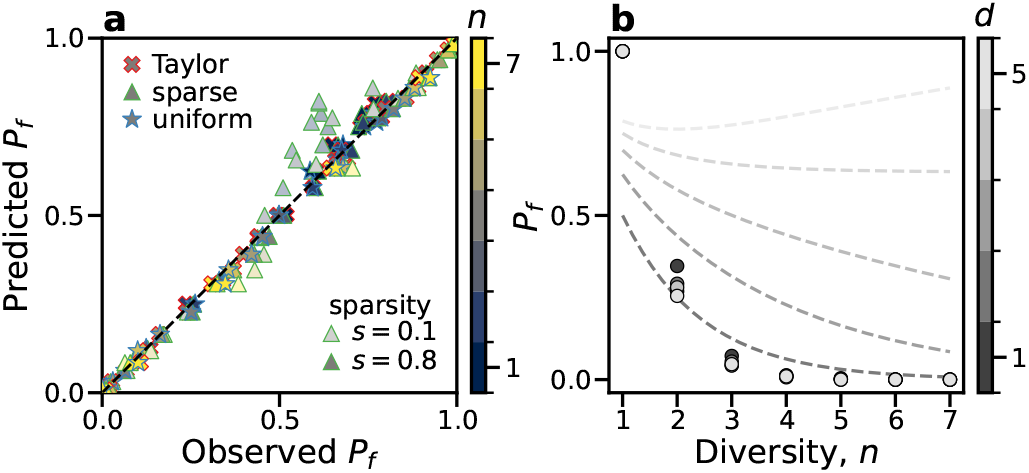
Universality and the role of sign symmetry. **(a)** Numerical estimate of *P*_*f*_ across the (*n, d*) grid versus the KSS independence approximation of Eq. (4) for the Taylor (cross), sparse (triangle), and uniform (star) ensembles. Color encodes species count *n*; for the sparse markers, lightness encodes sparsity *s*. The dashed line is *y* = *x*. **(b)** Transition vanishes under broken sign symmetry. Same as in Fig. 1**a**, but for the truncated (purely competitive) ensemble.

To probe whether the transitions survives the dense all-to-all property of the KSS ensemble, we impose sparsity on each coefficient on a monomial of total degree |*l*| ≥ 3 by retaining it with probability *s* and zeroing it otherwise, while pairwise couplings and growth rates (|*l*| ≤ 2) are kept intact. Across *n* and *d, P*_*f*_ is essentially independent of *s*. The sparse markers in Fig. 2**a** (triangles) are well-aligned with theory even with sparsity as low as *s* = 0.1.

Next, we relax the assumption of Gaussian entries by sampling interactions from a Uniform distribution with variance decaying according to Eq. 5 (Fig. 2**a**, stars). Again, we observe no change, suggesting that the phase transition is universal (that is, independent of the distribution, as long as it is symmetric around 0).

Finally, we break the sign-symmetry of the entries by restricting all interaction coefficients (|*l*| ≥ 2) to the negative half-line while restricting the growth rates to be positive, yielding a purely competitive ensemble. Unlike the three previous relaxations, the transition does not survive: *P*_*f*_ decays monotonically with *n* at every *d* we tested, and the *d* = 4 crossover vanishes (Fig. 2**b**. The transition therefore depends on the sign-symmetry of the interaction coefficients, but not on the variance decay, sparsity, or marginal distribution.

## DISCUSSION

The transition at *d* = 4 is a counting race between two exponentials. Bézout’s theorem caps the number of complex equilibria at *d*^*n*^; orthogonal invariance of any sign-symmetric coefficient distribution sends each real root into one of 2^*n*^ orthants with equal weight. The expected number of feasible roots is therefore the real-root count 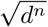 divided by the orthant count 2^*n*^, and these two exponentials cross at 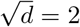 . As such, feasibility emerges whenever the rate at which a polynomial system produces real solutions overtakes the rate at which a symmetric distribution scatters them across orthants.

The transition survives modifications relevant to ecological realism. Namely, (1) imposing a variance structure resembling the structure emerging from Taylor expansion of a nonlinear system; (2) sparsifying the higher-order coefficients; and (3) sampling interaction terms from a Uniform (rather than Gaussian, suggesting universality) distribution. Emergent feasibility dis appeared, however, when the distributions were not symmetric around 0. Real ecological parameters are not generically sign-symmetric, with dominantly competitive/facilitative skewing the distribution toward negative or positive. If multiplicity of equilibria is what drives *P*_*f*_ → 1 in real ecosystems, it does so where interactions signs are mixed.

Feasibility is necessary for coexistence but not sufficient: a feasible equilibrium must also be dynamically stable. Recent results for pairwise competitive communities guarantees stability if feasibility is achieved [61]. Here we report the opposite: once stability is achieved, feasibility is guaranteed beyond a critical level of non-linearity. Characterizing the stability of feasible high-*d* equilibria would be a the natural next problem (see SI Sec. XII for preliminary results).

Enough nonlinearity, together with statistical sign symmetry suffices to make feasibility generic at high diversity; no ecological mechanism need be invoked to recover it such as self-regulation, intrinsic growth-rate sign structure, predator-prey architecture, modularity or niche partitioning [2, 5, 6, 8, 61]. This emergent feasibility is consistent with the notion of coexistence being the rule rather than the exception in ecological communities [44].

## METHODS

### Augmentation and polynomial homogenization

We collect the growth rate, pairwise couplings, and higher-order coefficients of species *i* in Eq. (1) into a single order-*d* tensor *T* ^(*i*)^ and homogenize. Promoting the environment to an auxiliary (*n*+1)-th coordinate of fixed value 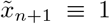 and writing 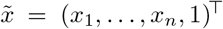, the right-hand side of Eq. (1) becomes the degree-*d* form

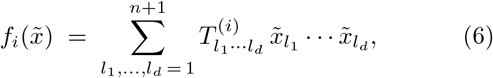

*n* homogeneous polynomials in the *n* + 1 variables 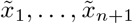, all of common degree *d*. A monomial with all *d* indices equal to *n* + 1 reproduces the growth rate *r*_*i*_; one with a single index *j* ≠ *n*+1 the pairwise term *A*_*ij*_*x*_*j*_; two the three-way term *B*_*ijk*_*x*_*j*_*x*_*k*_; and so on [52, 55– 57, 62].

Sampling every entry of *T* ^(*i*)^ as an independent mean-zero, unit-variance Gaussian places the system in the KSS ensemble, and indeed corresponds to ecological coefficients *r*_*i*_, *A*_*ij*_, etc being sampled from mean-zero Gaussians with variance 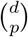 . The full correspondence is given in SI Secs. II–IV and VI.

### Root count and roots at infinity

By Bézout’s theorem, a generic degree-*d* system of *n* homogeneous polynomials in *n* + 1 variables has exactly *d*^*n*^ complex roots, with probability one over the random coefficients. The expected number of these that are real is E[*N*_ℝ_] = *d*^*n/*2^, the Kostlan-Shub-Smale-Edelman identity [53]. The ecologically relevant roots are those on the affine chart 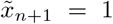, i.e. in ℝ^*n*^; any remaining roots would sit at infinite abundance, 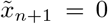 . Setting 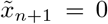 collapses the system to *n* homogeneous polynomials in only *n* variables, an overdetermined system that generically has no solution, so with probability one none of the *d*^*n*^ roots escapes to infinity and the count carries over to ℝ^*n*^. A self-contained treatment in projective space, the integral-geometry framework underlying the identity, and its extension to other random ensembles are given in SI Secs. I and IX–XI.

### Symmetry argument

The expected count of feasible equilibria in Eq. (3) differs from the real-root count of Eq. (2) by a factor 2^*−n*^. This is the statement that, in expectation, real roots are distributed uniformly across orthants. We show this.

Index the 2^*n*^ orthants of ℝ^*n*^ by *o*, and let *N*_*o*_ count the real roots falling in orthant *o*, so that *N*_ℝ_ = Σ_*o*_ *N*_*o*_ and, by linearity of expectation, E [*N*_ℝ_] = Σ_*o*_ E[*N*_*o*_]. We claim every term in this sum is equal. Consider the reflection (*x*_1_, …, *x*_*n*_) *→* (−*x*_1_, *x*_2_, …, *x*_*n*_). Applied to a KSS system, it flips the sign of every coefficient attached to a monomial of odd degree in *x*_1_; since these coefficients are sampled from a centered Gaussian, the reflected system has the same law as the original. The map is a bijection between the orthant with *x*_1_ *>* 0 and its *x*_1_ *<* 0 counterpart (remaining signs fixed), so the two carry equal expected root counts. Composing such reflections coordinate by coordinate connects any orthant to any other, hence E[*N*_*o*_] is independent of *o* and

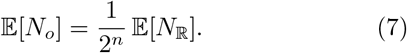

Taking *o* to be the positive orthant gives E[*N*_+_] = 2^*−n*^ E [*N*_ℝ_], and substituting Eq. (2) recovers Eq. (3). The argument uses only that the coefficient law is symmetric about zero coordinate-wise; it is otherwise distribution-free.

### Upper bound on the probability of feasibility

Let *N*_ℝ_ count the real zeros of the polynomial system and *N*_+_ the subset of those zeros lying in the positive orthant. Feasibility is the event *{N*_+_ *≥* 1*}*, so

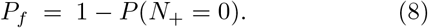

By orthogonal invariance of the KSS measure, every real root is equally likely to land in any of the 2^*n*^ orthants of ℝ^*n*^. Treating the orthant assignments of distinct real roots as independent (a working approximation that ignores correlations between sign patterns), conditional on *N*_ℝ_ = *j* the probability that none of the *j* real roots is positive is (1 *−* 2^*−n*^)^*j*^. Averaging over the law of *N*_ℝ_,

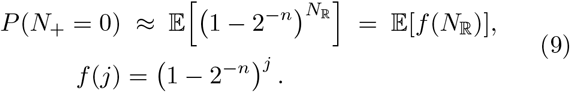

The function *f* is convex on [0, ∞ ): its second derivative is *f* (*j*) log^2^(1 − 2^*−n*^) *>* 0. Jensen’s inequality for convex *f* then gives

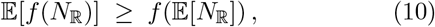

and combining (8)–(10) yields an upper bound on the approximated probability of feasibility,

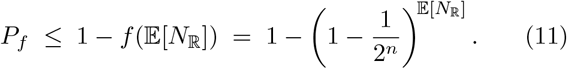

Substituting the identity E[*N*_ℝ_] = *d*^*n/*2^ from Eq. (2) into recovers the approximation (4) used in the main text.

### Variance schedule for the Taylor ensemble

Write the per-capita growth rate *f*_*i*_ : ℝ^*n*^ → ℝ as a degree-*d* Taylor polynomial around the origin,

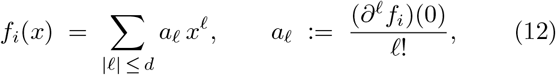

where *l* = (*l*_1_, …, *l*_*n*_) is a multi-index of total degree |*l*| = Σ_*j*_ *l*_*j*_, *l*! := *l*_1_! *· · · l*_*n*_!, and 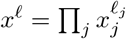 .

We postulate that the partial derivatives at the origin are i.i.d. standard Gaussians, (*∂*^*l*^*f*_*i*_)(0) *∼ N* (0, 1), independently across *i* and *l*. This is the smooth-function analogue of the centered Gaussian prior used for the KSS baseline: a maximally agnostic, sign-symmetric law on the derivatives of *f*_*i*_ at the origin. Because *a*_*l*_ is a deterministic rescaling of (*∂*^*l*^*f*_*i*_)(0) by the constant 1*/l*!, Var(*cX*) = *c*^2^Var(*X*) gives

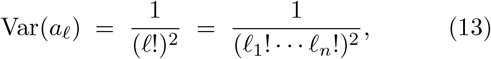

recovering Eq. (5). The factorial suppression encodes the smoothness prior: higher-order partials are penalized geometrically through *l*!, so the typical magnitude of a degree-|*l*| contribution to *f*_*i*_ decays as 1*/l*! at fixed point in parameter space. Sign symmetry of the coefficient law is preserved, so the orthant-uniformity argument of Methods B carries over.

## Acknowledgments

P.L.A. acknowledges E. Alastrue De Asenjo, C. Améndola, M. Bogan, D. Costa, J. Hoskins, S. Kuhen, S. Kundu, K. Lee, P. Lemos-Costa, S. Li, A. Liaghat, J. Lindberg, M. Loschi, M. Pascual, J. I. Rodriguez, S. Sheik, M. Silber, A. Skwara, M.-Ş. Sorea,L. Souto Ferreira, H.-Y. Tsai, T. Wootton, and P. Y. Yu for helpful discussions. P.B. acknowledges A. Lerario for for helpful discussions.

## Author contributions

P.L.A. designed the study and performed the research, with advice from A.S. S.A. and P.B.; P.L.A. wrote the manuscript; P.L.A. developed the code; P.L.A, A.S., P.B., and S.A. performed the derivations. All authors edited the manuscript, discussed and interpreted the results.

## Code and Data Availability

All code and data necessary to reproduce the paper can be found at https://github.com/pablolich/emergent_feasibility

