## supplementary information for "Emergent feasibility in random ecological systems with higher-order interactions"

### Supplementary Material

Pablo Lechón-Alonso,[1,*] Alexander Strang,[2] Paul Breiding,[3] and Stefano Allesina[1]

[1]*Department of Ecology & Evolution, University of Chicago,*

*1101 E. 57th Street, Chicago, IL 60637, USA*

[2]*Department of Statistics, University of California,*

*Berkeley, 110 Sproul Hall, Berkeley, CA, 94704, USA*

[3]*University of Osnabrück, Albrechtstr. 28a, 49076 Osnabrück, Germany*

(Dated: June 8, 2026)

*Reading guide.* This supplement is arranged so that a reader after one specific result can go straight to it. Section I fixes notation and recalls the projective-geometry background for counting equilibria. Sections II–IV build the random GLV model with higher-order interactions in three increasingly explicit forms—compact tensor (Sec. II), traditional (Sec. III), and polynomial (Sec. IV). Section V introduces the Kostlan–Shub–Smale (KSS) random polynomial system and its known root counts, and Sec. VI shows that the random GLV model is exactly such a system. Sections VII–VIII treat the ecologically central quantity, the probability of feasibility $P_f$: first a numerical and analytical estimate (Sec. VII), then an exact, assumption-free upper bound from linear programming (Sec. VIII). Sections IX–X recast everything in the language of integral geometry (Sec. IX) and extend the counts to general orthogonally invariant ensembles (Sec. X), and treat ensembles that are not orthogonally invariant (Sec. XI). Section XII shows that the Jacobian of a KSS ensemble is a Gaussian matrix. Finally, an Appendix reports numerical relaxations of the modeling assumptions. A reader interested only in feasibility can skip directly to Sections VII–VIII; one interested in the underlying mathematics will find it concentrated in Sections IX–X.

#### I. PRELIMINARIES

##### A. Notation

Here we fix the notation used throughout and review the geometric setting—projective space—needed to count the equilibria of polynomial systems.

Throughout this material we will be dealing with scalars, matrices, and tensors of arbitrary order. Scalars are denoted by non-capitalized letters with subscripts. For example, the abundance of population $i$ is a scalar, denoted by $x_i$. Vectors are obtained by dropping the index. For example, the abundance vector for the populations in a community is $x$. Matrices are denoted by capital letters. For example, the matrix with pairwise interactions of a community is denoted by $A$. Tensors of order greater than 2 are also denoted by capital letters. To avoid confusion, their dimension is stated when they are introduced. Slices of tensors are denoted by capital letters with subscripts. For example, if $B$ is a tensor of order 3 (a 3 dimensional array $3 \times 3 \times 3$), then $B_i$ is a slice of such tensor, that is, a $3 \times 3$ matrix, and $B_{ij}$ is a $3 \times 1$ column vector.

*

Multi-index notation will also be used throughout. As such, we will mark multi-indices in bold font, and indices in regular font. For example, the multi-index $\boldsymbol{j}$ containing $n$ different indices is denoted $\boldsymbol{j} = (j_1, \dots j_n)$.

Derivatives with respect to time are represented with dots, i.e. $\frac{dx}{dt} = \dot{x}$

##### B. Projective Space

In algebraic geometry, projective space is the preferred space when studying zero sets of systems of polynomial equations. It is obtained by adding the *hyperplane at infinity* (1) to *affine space* $\mathbb{C}^n$ (or $\mathbb{R}^n$). The reason why projective space is preferred over affine space is that it is compact: zeros of polynomial systems cannot "escape" to infinity.

The formal definition of (complex) projective space $\mathbb{P}^n$ is as the set of lines through the origin in $\mathbb{C}^{n+1}$:

$$\mathbb{P}^n := (\mathbb{C}^{n+1} \setminus \{0\}) / \sim, \\ \text{where } x \sim y \;:\Leftrightarrow\; \exists \lambda \in \mathbb{C} \setminus \{0\} : x = \lambda y \quad .$$

A line in $\mathbb{C}^{n+1}$/a point in $\mathbb{P}^n$ can be described by coordinates $[x_1 : \cdots : x_{n+1}]$ of one point on the line. The colon notation underlines $[x_1 : \cdots : x_{n+1}] = [\lambda x_1 : \cdots : \lambda x_{n+1}]$ for every $\lambda \in \mathbb{C}$, $\lambda \neq 0$.

We have an embedding $\mathbb{C}^n \hookrightarrow \mathbb{P}^n$ given by $(x_1, \dots, x_n) \mapsto [x_1 : \cdots : x_n : 1]$. This shows we have a disjoint union

$$\mathbb{P}^n = \mathbb{C}^n \cup H \quad , \tag{1}$$

where $H = \{[x_1 : \cdots : x_{n+1}] \in \mathbb{P}^n \mid x_{n+1} = 0\}$ is the hyperplane at infinity.

Similarly, we define real projective space $\mathbb{RP}^n$ as the set of lines through the origin in $\mathbb{R}^{n+1}$. In the same way, $\mathbb{RP}^n$ is the disjoint union of $\mathbb{R}^n$ and the real hyperplane at infinity. To get some intuition about projective space, think about the cases $n = 1$ and $n = 2$. The real projective line $\mathbb{RP}^1$ is obtained by taking the real line $\mathbb{R}$ and adding a point at infinity; i.e., we connect the loose ends of $\mathbb{R}$ at infinity thus obtaining a circle. For the real projective plane $\mathbb{RP}^2$ we add a line at infinity to the plane $\mathbb{R}^2$. This line is obtained by adding a point at infinity for every direction in $\mathbb{R}^2$ (but opposite directions are identified).

Finally, we define a notion of volume on real projective space. We have a $2:1$ map $p : S^n \to \mathbb{RP}^n$ of the $n$-dimensional sphere $S^n$ in $\mathbb{R}^{n+1}$ onto $\mathbb{RP}^n$ defined by $p(x_1, \dots, x_{n+1}) = [x_1 : \cdots : x_{n+1}]$. As a subset of $\mathbb{R}^{n+1}$ the sphere carries the usual Lebesgue measure, which means we can measure volumes of (measurable) subsets of the sphere. If we have subset $U \subset \mathbb{RP}^n$ such that $p^{-1}(U)$ is $k$-dimensional and measurable, we define the $k$-dimensional projective volume to be

$$\mathrm{vol}_k(U) := \frac{\mathrm{vol}_k(p^{-1}(U))}{2} \quad .$$

For instsance,

$$\mathrm{vol}_n(\mathbb{RP}^n) := \frac{1}{2}\mathrm{vol}_n(S^n) = \frac{\pi^{(n+1)/2}}{\Gamma(\frac{n+1}{2})} \quad .$$

We can now place our system in this setting. Written homogeneously, its equilibria are points in $\mathbb{P}^n$, and Bézout's theorem guarantees that a generic degree-$d$ system of $n$ such polynomials has exactly $d^n$ complex roots in $\mathbb{P}^n$, with probability one over the random coefficients. The ecologically relevant roots are those on the affine chart $\tilde{x}_{n+1} = 1$, i.e. in $\mathbb{R}^n$; by the disjoint union (1) any remaining roots would lie on the hyperplane at infinity $\tilde{x}_{n+1} = 0$. Setting $\tilde{x}_{n+1} = 0$ reduces the system to $n$ homogeneous polynomials in only $n$ variables, an overdetermined system that generically has no solution in $\mathbb{P}^{n-1}$; hence, with probability one, none of the $d^n$ roots escapes to infinity. The complex count, and with it the expected real count $\mathbb{E}[N_{\mathbb{R}}] = d^{n/2}$ stated in Sec. V, therefore holds over $\mathbb{R}^n$.

With notation and the projective picture in hand, we now introduce the ecological model.

#### II. GLV WITH HOIS

Consider a GLV model representing the abundance dynamics of $n$ populations, which we write as

$$\dot{x_i} = x_i \sum_{j=1}^{n+1} T_{ij}\tilde{x}_i \quad , \tag{2}$$

where $T$ is a $n \times (n+1)$ matrix of coefficients, and $\tilde{x}$ is an $(n+1) \times 1$ vector where the first $n$ components are population abuncances, and the last component is the constant 1

$$T = \begin{pmatrix} A_{11} & \dots & A_{1n} & r_1 \\ \vdots & \ddots & \vdots & \vdots \\ A_{n1} & \dots & A_{nn} & r_n \end{pmatrix} \quad \tilde{x} = \begin{pmatrix} x_1 \\ \vdots \\ x_n \\ 1 \end{pmatrix}$$

Consider now the extension of model 2 to account for higher-order interactions. In the case of three-way HOIs, the model can be written as

$$\dot{x}_i = x_i \sum_{jk}^{n+1} T_{ijk} \tilde{x}_i \tilde{x}_k \quad . \tag{3}$$

For generalized HOIs of order $d$, this model reads

$$\dot{x}_i = x_i \sum_{j_1 \dots j_d}^{n+1} T_{ij_1 \dots j_d} \tilde{x}_{j_1} \dots \tilde{x}_{j_d} \quad . \tag{4}$$

We assume the simplest random parametrization of the tensor $T$. Each entry of $T$ is sampled independently from a standard Gaussian distribution, i.e., $T_{ij_1 \dots j_d} \sim N(0,1)$

Next we unpack this tensor to recover the familiar GLV equations and to see how the interaction order rescales each coefficient.

#### III. TRADITIONAL REPRESETATION OF GLV WITH HOIS

Expanding the compact model term by term, we recover the traditional GLV equations and show that a coefficient of order $p$ is multiplied by the binomial factor $\binom{d}{p}$.

Note that if we expand the matrix multiplication of model 2, we can re-write the GLV in its traditional form, that is:

$$\begin{aligned} \dot{x}_i &= x_i \left( \sum_{j=1}^{n} T_{ij} \tilde{x}_j + T_{in+1} \tilde{x}_{n+1} \right) \\ &= x_i \left( \sum_{j=1}^{n} A_{ij} x_j + r_i \right) \quad . \end{aligned} \tag{5}$$

This tensor, partitioned in its inner and outer layers is represented in the top row of Fig. 1. The variances of the coefficients $A_{ij}$ and $r_i$ are also plotted next to it.

Similarly, if we expand Eq. 3 by leaving out the terms where any of the $x_i$ reaches the $n+1$

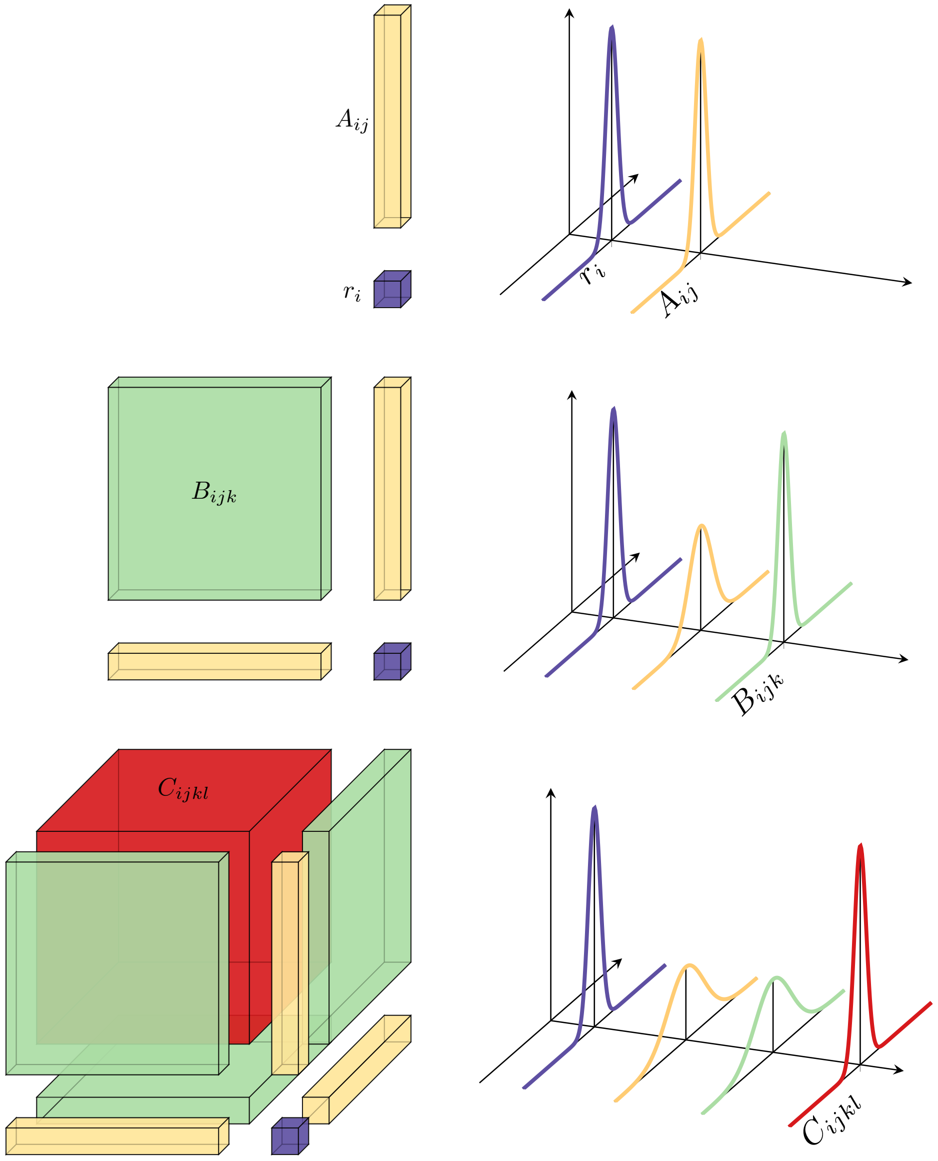


FIG. 1. The $i^{th}$ slice of the tensor $T$ ($T_i$) for $d = 1$ (pairwise interactions, top), $d = 2$ (three-way interactions, middle), and $d = 3$ (four-way interactions, bottom). For three-way interactions ($d = 2$), the matrix $T_i$ is composed of one sub-matrix, $B_i$, two vectors, $A_{ij}$, and one scalar $r_i$. When we perform the summation in Eq. 4 green matrix $B_i$ will give rise to quadratic terms of the form $B_{ijk}x_jx_k$. The yellow vectors $A_i$ are on the nth row and column of the matrix $T_i$. As such, they will give rise linear terms of the form $A_{ij}x_jx_n$ and $A_{ij}x_nx_j$, which are linear because $x_n = 1$. Finally, the purple scalar $r_i$ will give rise to 0th order terms of the form $r_ix_nx_n$. In the case of $d = 3$, $T_i$ is now a tensor, but the same reasoning as above applies. The only difference arises in the variance structure, due to the different geometries of dimensions 2, and 3. For $d = 2$, the variances of $B_{ijk}$ and $r_i$ will be 1, but those of $A_{ij}$ will be 2. For $d = 3$, the variances of $C_{ijkl}$ and $r_i$ will be 1, while those of $A_{ij}$ and $B_{ijk}$ will be 3.

component, we get

$$\begin{aligned}
\dot{x}_i &= x_i \left( \sum_{j,k=1}^{n} T_{ijk}\tilde{x}_j\tilde{x}_k + \sum_{k=1}^{n} T_{in+1k}\tilde{x}_{n+1}\tilde{x}_k + \sum_{j=1}^{n} T_{ijn+1}\tilde{x}_j\tilde{x}_{n+1} + T_{in+1n+1}\tilde{x}_{n+1}\tilde{x}_{n+1} \right) \\
&= x_i \left( \sum_{j,k+1}^{n} B_{ijk}x_jx_k + \sum_{k=1}^{n} A_{ik=1}x_k + \sum_{j=1}^{n} A_{ij}x_j + r_i \right) \\
&= x_i \left( \sum_{j,k=1}^{n} B_{ijk}x_jx_k + 2\sum_{k=1}^{n} A_{ik}x_k + r_i \right) \qquad .
\end{aligned}$$

The graphical intuition for this case is plotted in the second row of Fig. 1. The left panel shows that in the $d = 2$ case the tensor slice $T_i$ has two one-dimensional edges (yellow columns). As such, when we add them together, the variance doubles, as seen in the right panel.

For the 4-way interaction model, when we separate the outer layer and the inner one, we also arrive to a GLV with generalized HOIs of the form

$$\begin{aligned}
\frac{\dot{x}_i}{x_i} = & \sum_{j,k,l}^{n} T_{ijkl}\tilde{x}_j\tilde{x}_k\tilde{x}_l \\
& + \sum_{k,l=1}^{n} T_{in+1kl}\tilde{x}_{n+1}\tilde{x}_k\tilde{x}_l + \sum_{k,l=1}^{n} T_{ijn+1l}\tilde{x}_j\tilde{x}_{n+1}\tilde{x}_l + \sum_{k,l=1}^{n} T_{ijkn+1}\tilde{x}_j\tilde{x}_k\tilde{x}_{n+1} \\
& + \sum_{l=1}^{n} T_{in+1n+1l}\tilde{x}_{n+1}\tilde{x}_{n+1}\tilde{x}_l + \sum_{k=1}^{n} T_{in+1kn+1}\tilde{x}_{n+1}\tilde{x}_k\tilde{x}_{n+1} + \sum_{j=1}^{n} T_{ijn+1n+1}\tilde{x}_j\tilde{x}_{n+1}\tilde{x}_{n+1} \\
& + T_{in+1n+1n+1}\tilde{x}_{n+1}\tilde{x}_{n+1}\tilde{x}_{n+1} \\
= & \sum_{j,k,l=1}^{n} C_{ijkl}x_jx_kx_l \\
& + \sum_{k,l=1}^{n} B_{ikl}x_kx_l + \sum_{j,l=1}^{n} B_{ijl}x_jx_l + \sum_{j,k=1}^{n} B_{ijk}x_jx_k \\
& + \sum_{j} A_{ij}x_j + \sum_{k} A_{ik}x_k + \sum_{l} A_{il}x_l + r_i \\
= & \, r_i + 3\sum_{k} A_{ik}x_k + 3\sum_{j,k=1}^{n} B_{ijk}x_jx_k + \sum_{j,k,l=1}^{n} C_{ijkl}x_jx_kx_l \qquad .
\end{aligned}$$

The graphical representation of this case is plotted in the third panel of Fig. 1.

In these four examples we have seen that depending on the order of the interactions in the system, each coefficient is scaled differently. By induction, for a GLV with HOIs of order $d$, we can write the general form of the scalar multiplying coefficient of order $p$ (where $p \leq d$) as

$$c_p^{(d)} = \binom{d}{p} \quad .$$

Since each coefficient has unit variance, we can absorb it into the coefficient, increasing the variance of the distribution from which it is sampled. As such, we can write per-capita growth-rate of the GLV with generalized HOIs as

$$\frac{\dot{x}_i}{x_i} = F(x_i, ..., x_n) = \Gamma_i^{(d)} + \sum_{j_1} \Gamma_{ij_1}^{(d)} x_{j_1} + \cdots + \sum_{j_1 \cdots j_p} \Gamma_{j_1 \cdots j_p}^{(d)} x_{j_1} \ldots x_{j_p} + \cdots + \sum_{j_1 \cdots j_d} \Gamma_{ij_1 \cdots j_d}^{(d)} x_{j_1} \ldots x_{j_p} \quad (6)$$

where the coefficients $\Gamma_{ij_1 \cdots j_p}^{(d)}$ are sampled from a uniform distribution with mean 0 and variance given by

$$\mathrm{Var}[\Gamma_{ij_1 \cdots j_p}^{(d)}] = \binom{d}{p}$$

Having tracked how the interaction order scales each coefficient, we now group terms by monomial to reach the polynomial form, where a finer variance structure appears.

#### IV. POLYNOMIAL REPRESENTATION OF GLV WITH HOIS

As seen in the previous section, the coefficients of the GLV model when expressed in its traditional form have a special variance structure. Here we perform one more step by grouping together coefficients associated with the same monomials. This last step expresses the GLV model in its polynomial form, that is, an expression where coefficients have been group according to monomials of the same order/configuration. In the case of interactions of order 0 and 1, given by Eq. 5, the polynomial form of the model is the same as the traditional form, since $r_i$ is the only coefficient of order 0 in equation $i$, and $A_{ij}$ is the only coefficient of order 1 for equation $i$ and population $j$ per population $j$. For three-way coefficients however, the correspondence is not unique. For example, monomials of the form $x_j x_k$, with $j \neq k$ will appear twice; once multiplying the coefficient $B_{ijk}$ and once multiplying the coefficient $B_{ikj}$. As such, we can group them as

$$B_{ijk} x_j x_k + B_{ikj} x_k x_j = (B_{ijk} + B_{ikj}) x_j x_k = a_{11}^{(i)} x_j x_k \quad .$$

On the contrary, the monomial $x_j^2$ will only appear once, associated with the coefficient $B_{ijj}$. As such, the variance of the polynomial coefficients associated with monomials of degree 2 of the example above are

$$\text{Var}[a^{(i)}_{|\boldsymbol{l}|=2}] = \begin{cases} \text{Var}[a^{(i)}_{11}] = 2 \\ \text{Var}[a^{(i)}_{20}] = 1 \end{cases} ,$$

where $\boldsymbol{l}$ is a set of $n$ natural numbers indicating the exponent of each variable in the monomial associated with the coefficient in the ith equation $a^{(i)}_{\boldsymbol{l}}$. $|\boldsymbol{l}|$ is the sum of the elements in the set, which is always 2 for three-way interactions. In the case of four-way interactions, we would have that monomials of the form $x_j x_k x_l$, where $j \neq k \neq l$, would appear three times

$$\begin{aligned} C_{ijkl}x_jx_kx_l + C_{iljk}x_lx_jx_k + C_{iklj}x_kx_lx_j + C_{ijlk}x_jx_lx_k + C_{ikjl}x_kx_jx_l + C_{ilkj}x_lx_kx_j = \\ (C_{ijkl} + C_{iljk} + C_{iklj} + C_{ijlk} + C_{ikjl} + C_{ilkj})x_jx_kx_l = \\ a^{(i)}_{111}x_jx_kx_l \end{aligned}$$

On the contrary, when the three indices take the same value, the corresponding monomial $x_j^3$ would only appear once

$$C_{ijjj}x_j^3 = a^{(i)}_{300}x_j^3 \quad .$$

Alternatively, if two coefficients take the same value but one is different, we get

$$(C_{ijjk} + C_{ijkj} + C_{ikjj}) = a^{(i)}_{210}x_j^2x_k \quad .$$

As such, the variance of the polynomial coefficients associated with monomials of degree 3 we considered in this example are

$$\text{Var}[a^{(i)}_{|\boldsymbol{l}|=3}] = \begin{cases} \text{Var}[a^{(i)}_{111}] = 6 \\ \text{Var}[a^{(i)}_{210}] = 3 \\ \text{Var}[a^{(i)}_{300}] = 1 \end{cases} .$$

The examples above show that the variance of the polynomial coefficient depends not only on the degree of its corresponding monomial (as the in the previous section describing the variance structure of the GLV coefficients), but also on its configuration. That is, on how the total degree is distributed among the variables. In general, given monomial of degree $p \leq d$ with configuration $\boldsymbol{l}$, to compute its variance we need to count the number of ways of arranging $p$ distinct objects into $n$ distinct bins, with $l_1$ objects in the first bin, $l_2$ objects in the second bin, and so on. This is precisely the combinatorial interpretation of the multinomial coefficient. Thus, expression for the variance of the polynomial coefficients of the GLV model has the form

$$\mathrm{Var}(a_{\boldsymbol{l}}^{(i)}) = \binom{d}{\boldsymbol{l}} = \frac{d!}{l_1! \dots l_n!(d-|\boldsymbol{l}|)!} \quad . \tag{7}$$

This polynomial structure is precisely that of the Kostlan–Shub–Smale ensemble, which we define next.

#### V. KSS SYSTEM OF RANDOM POLYNOMIALS

We introduce the Kostlan–Shub–Smale (KSS) ensemble of random polynomials and recall its known root counts: the expected number of real roots, the vanishing of the normalized variance, and the resulting count of positive roots.

Consider a system $\boldsymbol{P}$ of $n$ polynomials with common degree $d > 1$, on $n$ variables. The $i^{th}$ equation of such system can be written as

$$P_i(x_1, \dots, x_n) = \sum_{|\boldsymbol{l}| \le d} a_{\boldsymbol{l}}^{(i)} \boldsymbol{x}^{\boldsymbol{l}} \quad , \tag{8}$$

where

1. $\boldsymbol{l} = \{l_1, \dots, l_n\} \in \mathbb{N}^{[0,n]}$, and $|\boldsymbol{l}| = \sum_{k=1}^{n} l_k$,
2. $a_{\boldsymbol{l}}^{(i)} = a_{l_1,\dots,l_n}^{(i)} \in \mathbb{R}$, $|\boldsymbol{l}| \le d$,
3. $\boldsymbol{x} = (x_1, \dots, x_n)$ and $\boldsymbol{x}^{\boldsymbol{l}} = \prod_{k=1}^{n} x_k^{l_k}$.

We say that the system $\boldsymbol{P}$ has the Kostlan-Shub-Smale (KSS) distribution [1, 13] if the coefficients $a_{\boldsymbol{l}}^{(i)}$ are i.i.d. random variables with mean 0 and variance

$$\mathrm{Var}(a_{\boldsymbol{l}}^{(i)}) = \binom{d}{\boldsymbol{l}} = \frac{d!}{l_1! \dots l_n!(d-|\boldsymbol{l}|)!} \quad . \tag{9}$$

Importantly, for this class of systems of polynomials, it has been shown [12] that the expected number of real roots is

$$\mathbb{E}[N_{\mathbb{R}}] = \sqrt{d^n} \tag{10}$$

Additionally, [14] showed that the variance of the normalized number of real roots vanishes in the for $d \ge 3$ in the limit of $n \to \infty$, that is

$$\lim_{n\to\infty} \mathrm{Var}\left[\frac{N_{\mathbb{R}}}{\sqrt{d^n}}\right] = 0 \quad . \tag{11}$$

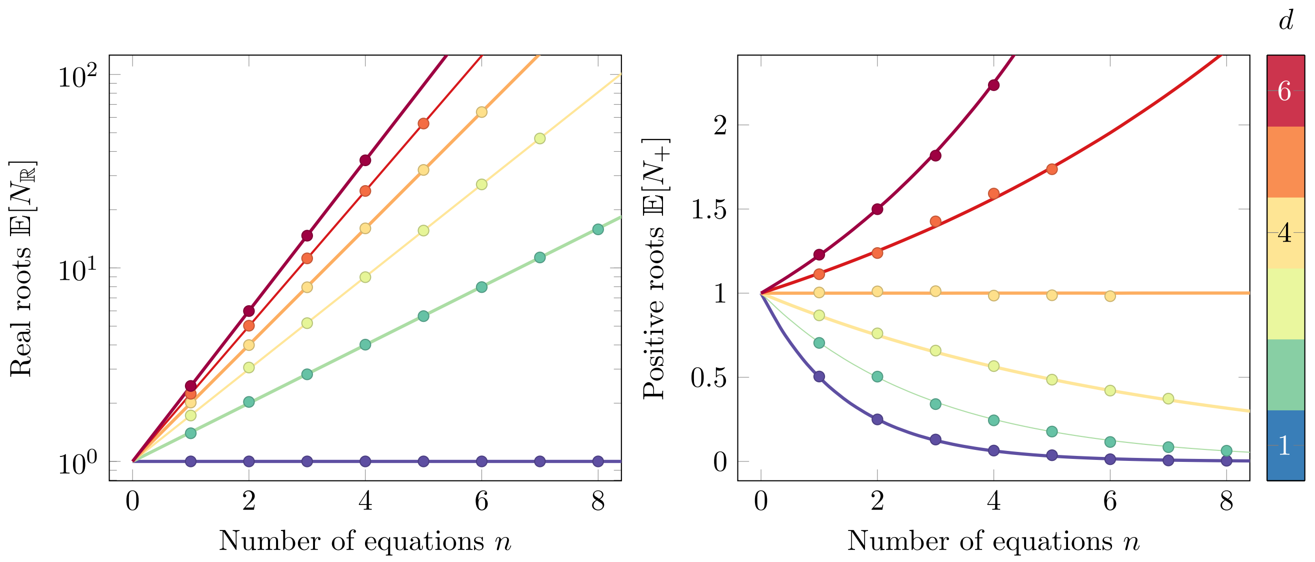


FIG. 2. **Expected number of real roots and positive roots.** On the left panel we plot (semilog scale) the expected number of real roots (Eq. 10) for different values of $d$, and overlay the average value of real roots obtained from simulations of the KSS polynomial system. On the right panel we plot the expected number of positive solutions. Importantly, this number grows with $n$ for $d > 4$

These two results can be easily cast into statements about the number of positive roots. To do so we rely in the fact that the coefficients of $P_i$ are sampled from a distribution centered at zero. As such, the sign of each component of the roots are not biased in any direction. Thus, the asymptotic behaviour of the variance of the positive roots will remain, and, by symmetry, the expected number of positive roots is just [4, 10]

$$\mathbb{E}[N_+] = \frac{1}{2^n}\mathbb{E}[N_{\mathbb{R}}] = \left(\frac{\sqrt{d}}{2}\right)^n \tag{12}$$

Numerical simulations of the system in Eq. 8 (Fig. 2, dots) using homotopy continuation [2, 8], track well both of the theoretical curves (Eqs. 10 and 12). Importantly, one can see from Eq. 12 that the expected number of positive roots becomes 1 for $d = 4$, and grows with $n$ for $d > 4$. In the next section, we establish the connection between this system and the GLV model. Thereafter, we make use of such connection to study the ecological consequences of the divergence of the number of positive roots.

These statements hold for any KSS system; we now show that the random GLV model defined above is indeed the KSS system, so all of them transfer to it.

#### VI. FROM KSS TO GLV

Here we show that the system of polynomials arising when solving for feasible equilibria of Eq. 2 is precisely the KSS system of random polynomials when the coefficients in $r, A, B, \ldots$ are sampled with mean 0, and appropriate variances. Recently, this equivalence has been shown for the replicator equation with higher order interactions [4].

First, we expand Eq. 8 as

$$\begin{aligned} P_i(x_1, \ldots x_n) &= \sum_{|\boldsymbol{l}| \leq d} a^{(i)}_{\boldsymbol{l}} \prod_{k=1}^{n} x_k^{l_k} \\ &= \sum_{|\boldsymbol{l}|=0} a^{(i)}_{0\ldots 0} + \sum_{|\boldsymbol{l}|=1} a^{(i)}_{l_1 \ldots l_n} \prod_{k=1}^{n} x_k^{l_k} + \cdots + \sum_{|\boldsymbol{l}|=d} a^{(i)}_{l_1 \ldots l_n} \prod_{k=1}^{n} x_k^{l_k} \quad . \end{aligned} \tag{13}$$

The first commonality between systems in Eqns. 13 and 6 is that they have the same total degree; that is $h = d$. We can now identify coefficients of Eq. 6 with the corresponding coefficients of Eq. 13 that multiply monomials of equal degree. For the first term in the sum of Eq. 13, that with $|\boldsymbol{l}| = 0$, there is only one monomial of degree 0; (that is, the scalar 1). As such, the corresponding coefficient in the GLV model (Eq. 6) is simply

$$r_i = a^{(i)}_{0\ldots 0}$$

For $r_i$ to be a coefficient of the KSS system, its variance should be that of Eq. 9

$$\mathrm{Var}[r_i] = \binom{d}{\boldsymbol{l}} = \frac{d!}{0!d!} = 1 \quad .$$

For the coefficients of the second term of the sum in Eq. 13 (those paired with monomials of degree $|\boldsymbol{l}| = 1$), the corresponding coefficients in the GLV model would be

$$\begin{aligned} \sum_{j_1} A_{ij_1} x_{j_1} &= \sum_{|\boldsymbol{l}|=1} a^{(i)}_{l_1 \ldots l_n} \prod_{k=1}^{n} x_k^{l_k} \\ A_{i1}x_1 + A_{i2}x_2 + \cdots + A_{in}x_n &= a^{(i)}_{10\ldots 0} x_1 + a^{(i)}_{01\ldots 0} x_2 + \ldots a^{(i)}_{00\ldots 1} x_n \\ A_{ij} &= a^{(i)}_{\boldsymbol{l}_j} \quad , \end{aligned}$$

where $\boldsymbol{l}_j$ denotes the multiindex of all zeros and a 1 in position $j$. As before, the variance of the $A_{ij}$ coefficients should be

$$\mathrm{Var}[A_{ij}] = \binom{d}{d-1} = d \quad ,$$

Next, we identify with the GLV coefficients associated with interactions of order 3, $B_{ijk}$, with the coefficients of monomials of degree $|\boldsymbol{l}| = 2$ in Eq. 13

$$\sum_{|\boldsymbol{l}|=2} a^{(i)}_{l_1 \ldots l_n} \prod_{k=1}^{n} x_k^{l_k} = \sum_{j_1 j_2} B_{ij_1 j_2} x_{j_1} x_{j_2}$$

$$\begin{aligned} B_{i11}x_1^2 + (B_{i12} + B_{i21})x_1 x_2 + \cdots +& \\ (B_{i1n} + B_{in1})x_1 x_n +& \\ (B_{i2n} + B_{in2})x_2 x_n + \cdots + B_{inn}x_n^2 &= a^{(i)}_{20\ldots0}x_1^2 + a^{(i)}_{11\ldots0}x_1 x_2 + a^{(i)}_{10\ldots1}x_1 x_n + \ldots \\ & \quad a^{(i)}_{10\ldots1}x_1 x_n + a^{(i)}_{01\ldots1}x_2 x_n + \cdots + a^{(i)}_{00\ldots2}x_n^2 \end{aligned}$$

The first thing to note from the above expression is that for $|\boldsymbol{l}| = 2$, we sometimes find more than one GLV parameter per KSS coefficient. In particular, when $j = k$ there correspondence is 1 to 1, but wehen $j \neq k$ it is 1 to 2. This is due to the commutative property of multiplication. Therefore, the variance of the $B_{ijk}$ coefficients appears to be index dependent. Let us, apply Eq. 9 to each case, we have that

$$\mathrm{Var}[B_{ijj}] = \mathrm{Var}[a^{(i)}_{0\ldots2\ldots0}] = \frac{1}{2}d(d-1)$$

and

$$\mathrm{Var}[B_{ijk}] + \mathrm{Var}[B_{ikj}] = \mathrm{Var}[a^{(i)}_{0\ldots1\ldots1\ldots0}]$$

Without loss of generality we assume the variances of coefficients with indices $\{j, k\}$ that are permutations of each other to be the same. Plugging in Eq. 9, we get

$$\mathrm{Var}[B_{i\sigma(jk)}] = \frac{1}{2}d(d-1) \quad ,$$

where $\sigma(jk)$ denotes permutations of the set $\{j, k\}$. Conveniently, the variance of the element $B_{ijk}$ turns out to not depend on its position on the tensor. As we will see next, this is not an artifact specific to order 2, but a general feature.

Lastly, we consider the general case of monomials of order $|\boldsymbol{l}| = p$

$$\sum_{|\boldsymbol{l}|=p} a^{(i)}_{l_1 \ldots l_n} \prod_{k=1}^{n} x_k^{l_k} = \sum_{j_1 j_2} T_{ij_1 \ldots j_p} x_{j_1} \ldots x_{j_p} \quad .$$

Due to the commutative property, identifying coefficients on the same monomials leads to the following equality for any given monomial of degree $p$

$$a^{(i)}_{l_1 \ldots l_n} = \sum_{\sigma(j_1 \ldots j_p)} T_{ij_1 \ldots j_p} \quad , \tag{14}$$

where the sum is taken over all the permutations of the multiset $\mathcal{S} = \{j_1 \dots j_p\}$. Taking the variance on both sides leads to

$$\mathrm{Var}[a^{(i)}_{l_1 \dots l_n}] = \mathrm{Var}\left[\sum_{\sigma(\mathcal{S})} T_{ij_1 \dots j_p}\right] = r\mathrm{Var}[T_{ij_1 \dots j_p}] \quad , \tag{15}$$

where the sum is not performed over different values of indices; that is, given a multiset of indices, $\mathcal{S}$, we sum over all the distinguishable permutations of such set $\sigma(\mathcal{S})$. There are a total of $r$ number of permutations of $\mathcal{S}$, and we have used that the variance of the sum of sum of variances since $T_{i\sigma(\mathcal{S})}$ are i.i.d.

Given a multiset of $p$ objects such that there are $p_1$ identical objects of type 1, $p_2$ identical objects of type 2... and $p_k$ identical objects of type $k$, the number of permutations of such multiset is

$$r = \frac{p!}{p_1! \cdots p_k!}$$

Note that since we are identifying coefficients, the numbers $p_1, \dots, p_k$ are indeed $l_1, \dots, l_k$. Making these substitutions and plugging in Eq. 9 into Eq. 15, we can solve for the variance of any coefficient of $T_{ij_1 \dots j_p}$ arriving to

$$\mathrm{Var}[T_{ij_1 \dots j_p}] = \frac{l_1! \cdots l_n!}{d!} \frac{d!}{l_1! \dots l_n!(d-p)!} = \frac{d!}{p!(d-p)!} = \binom{d}{p} \quad . \tag{16}$$

Surprisingly, this variance is only depends on the coefficient order, $p$, and the overall order of HOIs in the ecosystem, $d$ (see Fig. 3). Note that the largest variance occurs when the maximum size of the group of simultaneously interacting populations (the order of the interaction, $d+1$) is largest (top row of Fig. 3). Specifically, the maximum is reached when the size of the interacting group of populations (coefficient order, $p$) is half of the maximum allowed size, $p = \frac{1}{2}(d+1)$.

We have shown that, as long as the coefficients of the GLV model are sampled from a Gaussian distribution with mean 0 and variance given by Eq. 16, then it holds that

$$F_i(x_1, \dots, x_n) \equiv P_i(x_1, \dots x_n) \quad .$$

As such, all the results derived for the KSS system of random polynomials can be applied to the random GLV with higher-order interactions.

With the equivalence in place, every KSS result applies to our GLV model. We turn to the quantity of ecological interest, the probability of feasibility.

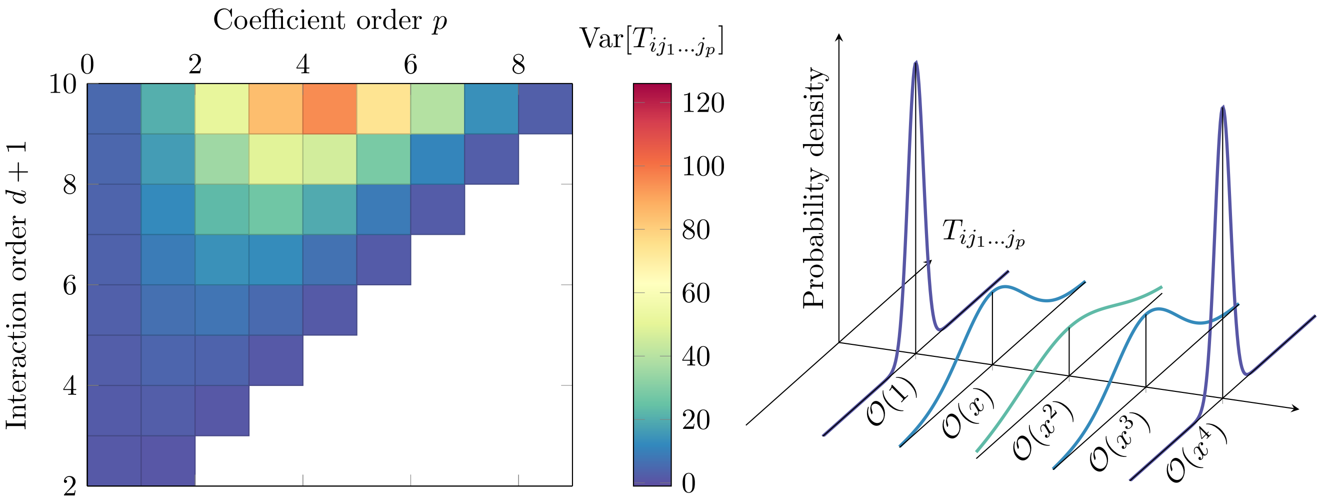


FIG. 3. **Variance of the GLV coefficients**. (left) We plot the variance obtained in Eq. 16 as a function of coefficient order $p$ (x-axis) and overall interaction order of the system $d$ (y-axis). For a given coefficient order $p$ (vertical stripes), its variance increases as the interaction order of the system increases. On the other hand, for a given interaction order $d$ (horizontal stripes), the variance of the coefficient reaches a maximum for intermediate values of its order. (right) For interaction order $d = 4$ we plot the probability distributions from which we sample the coefficients associated to each monomial. The extreme monomials, with powers 0 and 4 have coefficients sampled from normal distributions with the smallest variance. The the middle monomials, those with degree 2 (we label them generically as $x^2$) are sampled from a distribution with the largest variance.

#### VII. PROBABILITY OF FEASIBILITY

Real ecosystems exhibit only one equilibrium at a time. Thus, the ecologically relevant quantity is the probability of feasibility; $P_f$. This is the probability of having at least one solution with all positive components, which we compute as

$$P_f = \sum_{i=1}^{d^n} P(N_+ = i) = 1 - P(N_+ = 0) \quad , \tag{17}$$

that is, we sum over the probabilities of having any number of feasible solutions greater than 0 ($i \in [1, d^n]$). The probability distribution $P(N_+)$ has discrete support in the integers $\mathbb{Z} \in [0, d^n]$, and expected value given by Eq. 12 but is otherwise unknown for arbitrary d. As such, we can only approximate $P(N_+)$ given a sufficiently large ensemble of equilibria of random GLV systems

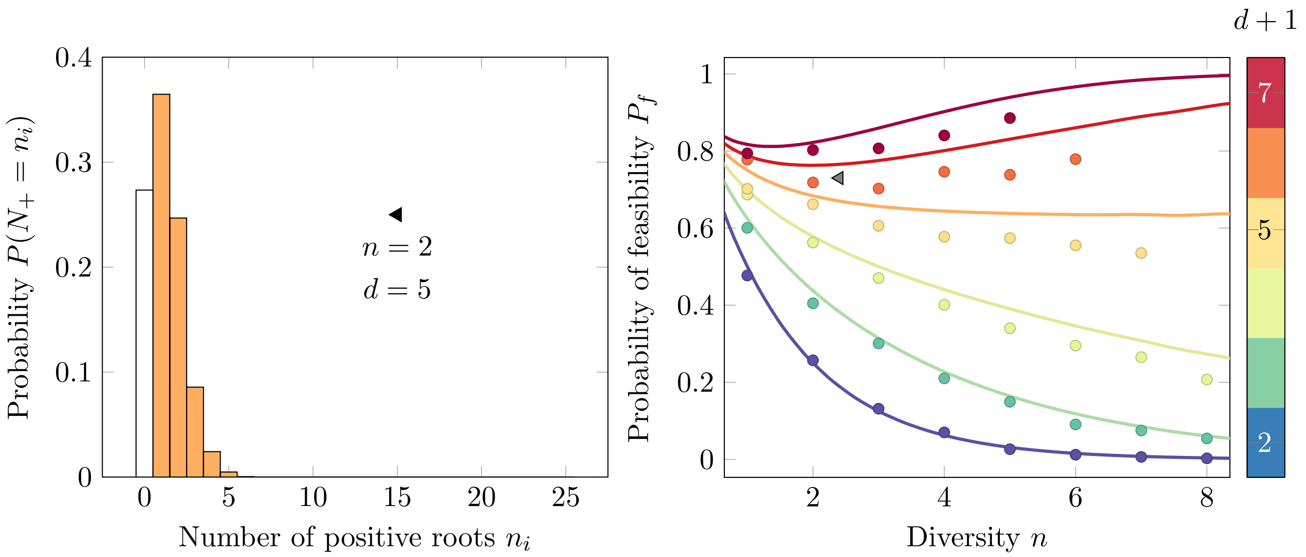


FIG. 4. Left: Numerical approximation of the probability distribution $P(N_+)$ for $d = 5$, and $n = 2$. The height of the $i^{th}$ bar is obtained by dividing the number of times that we observe $i$ positive solutions by the total number of carried simulations (in this case, 1000). Right: probability of feasibility ($P_f$, y-axis) as a function of diversity ($n$, x-axis) and interaction order ($d$, colors) for computationally tractable values of $n$, and $d$. $P_f$ decreases with $n$ for $d \leq 5$, but approaches 1 for $d > 5$. Points and lines are numerical and theoretical approximations, respectively. Black triangular marker connects the two plots: that is, adding the heights of the orange bars in the left yields the numerical approximation of $P_f$ (y-axis) plotted on the right.

obtained numerically. An example of this numerical approximation for $d = 5$ and $n = 2$ is shown in the histogram of Fig. 4. We then can compute $P_f$ for different values of $n$ and $d$ using Eq. 17. This would be equivalent to adding the coloured bars of the histogram in Fig. 4. In the right panel of Fig. 4, we show numerical estimations (points) of $P_f$ for computationally tractable values of $n$ and $d$. Furthermore, we can try to approximate $P_f$ analytically by subtracting the probability to have 0 positive roots from one, obtaining:

$$\begin{aligned} P_f &\approx 1 - (p_1 p_- + p_2 p_- p_- + \cdots + p_{d^n} p_-^{d^n}) \\ &= 1 - \sum_{n_i=0}^{d^n} p_i \left(1 - \frac{1}{2^n}\right)^{n_i} \\ &= 1 - \mathbb{E}[f(n_i)] \quad . \end{aligned} \tag{18}$$

Here, $p_i$ is the probability that system 6 has $n_i$ real roots. The function $f(n_i) = (1-2^{-n})^{n_i}$, is the probability that none of the $n_i$ roots are feasible. We sum over all possible number of real roots; from 0 (found when all roots are complex) to $d^n$ (when all roots are real). In Eq. 18 we have assumed independence in the probability of having no positive roots; that is, the probability that one root has does not have all positive components does not depend on the sign of the components of other roots. This assumption only works for $d = 1$, because the number of roots is 1. In fact, the case where $d = 1$ has been well characterized by [11] (blue line in Fig. 4). In this case, $d^n = 1$, and Eq. 18 reduces to

$$P_f = \frac{1}{2^n} \quad .$$

Note however, that this approximation is not exact for arbitrary $d$.

Since we know the first moment of the random variable $n_i$ (Eq. 10), we can use it to approximate $\mathbb{E}[f(i)]$ through a Taylor expansion around the mean

$$\begin{aligned} P_f &\approx 1 - \left( f(\bar{X}) + \frac{df}{dn_i}\Big|_{n_i=\bar{X}} \mathbb{E}[n_i - \bar{X}] + \mathcal{O}\left(E[(n_i-\bar{X})^2]\right) \right) \\ &\approx 1 - \left(f(\bar{X})\right) \\ &\approx 1 - \left(1 - \frac{1}{2^n}\right)^{d^{n/2}} \quad . \end{aligned} \tag{19}$$

Here we have truncated the expansion to the first order. This truncation becomes more accurate as $n$ increases, since the variance vanishes in the limit of high diversity (Eq. 11, [14]). As such, the expected value of the number of positive roots is the first-order statistical approximation of the random variable $N_+$. In Fig 4, we plot this approximation in solid lines. In Fig 5 (left panel) we show that the approximation is very good even for moderately high $n$, and thus we plot the heat map (right panel) of $P_f$ as a function of $d$ and $n$ obtained from Eq. 19. In this limit, the scenario that we would see is that the approximated $P_f$ converges to 0 if $d < 4$, or 1 if $d > 4$. Importantly, the first order approximation is an upper bound for $P_f$. We can see this using Jensen's inequality, which has the form

$$\psi\left(\sum_i p_i x_i\right) \leq \sum_i p_i \psi(x_i) \quad , \tag{20}$$

where $p_i \geq 0$, $\sum_i p_i = 1$, and $\psi$ is convex. In our case, $\psi(x_i) = f(n_i)$. This function is convex on $n_i$, since its second derivative

$$\frac{d^2 f}{dn^2} = f(n_i) \log^2\left(1-2^{-n}\right) > 0$$

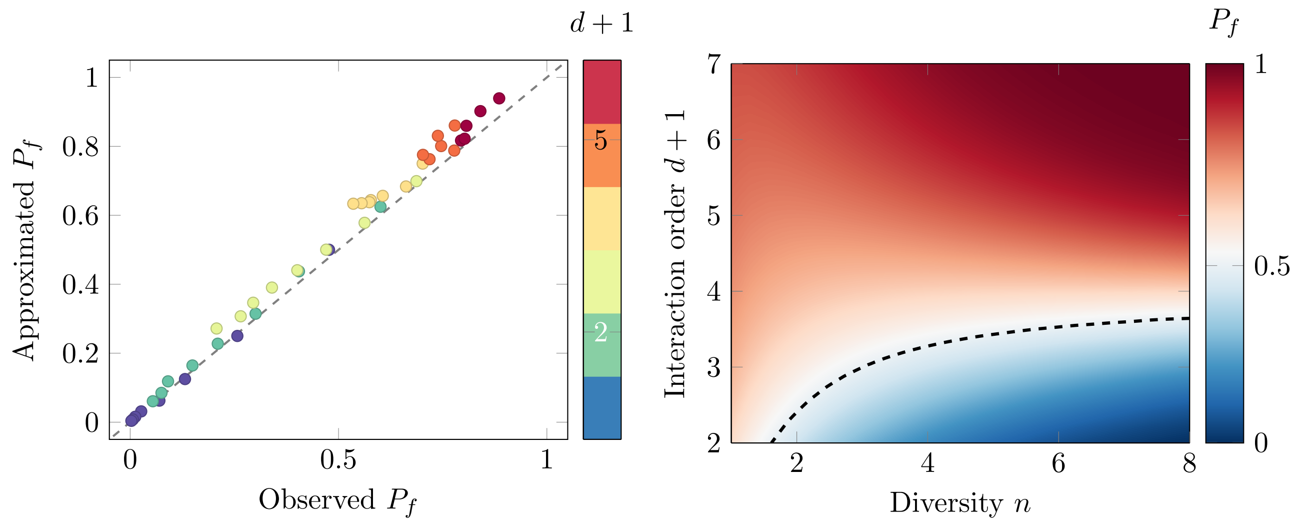


FIG. 5. (left) Approximated probability of feasibility versus the observed probability of feasibility in simulations. Different colors correspond to different values of interaction order $d$, and points within one color represent different values of $n$. The one to one line (dashed grey) is plotted for reference. (right) Heat map of the upper bound of $P_f$ (computed from Eq. 19) as a function of both $n$ and $d$. Level set where $P_f = 0.5$ is plotted as dashed line for reference.

is always positive. We apply this inequality to Eq. 18 by multiplying Eq. 20 by -1, and adding 1, arriving to

$$1 - \sum_i p_i f(n_i) \leq 1 - f\left(\sum_i p_i n_i\right)$$
$$P_f \leq 1 - f(\bar{X}) \quad . \tag{21}$$

This estimate rests on an independence assumption that is uncontrolled at finite diversity; we next replace it with an exact bound.

#### VIII. AN EXACT UPPER BOUND TO $P_f$

The analytical expression in Eq. 19 is an approximation based on the assumption that the signs of polynomial root are independent from each other. Here, we compute an exact upper bound to $P_f$ using linear programming.

$P_f$ is the probability to have at least one (but possibly more than one) positive equilibrium. To calculate this probability we have to add the probabilities to have $i$ positive roots, $p_i$ where $i \neq 0$

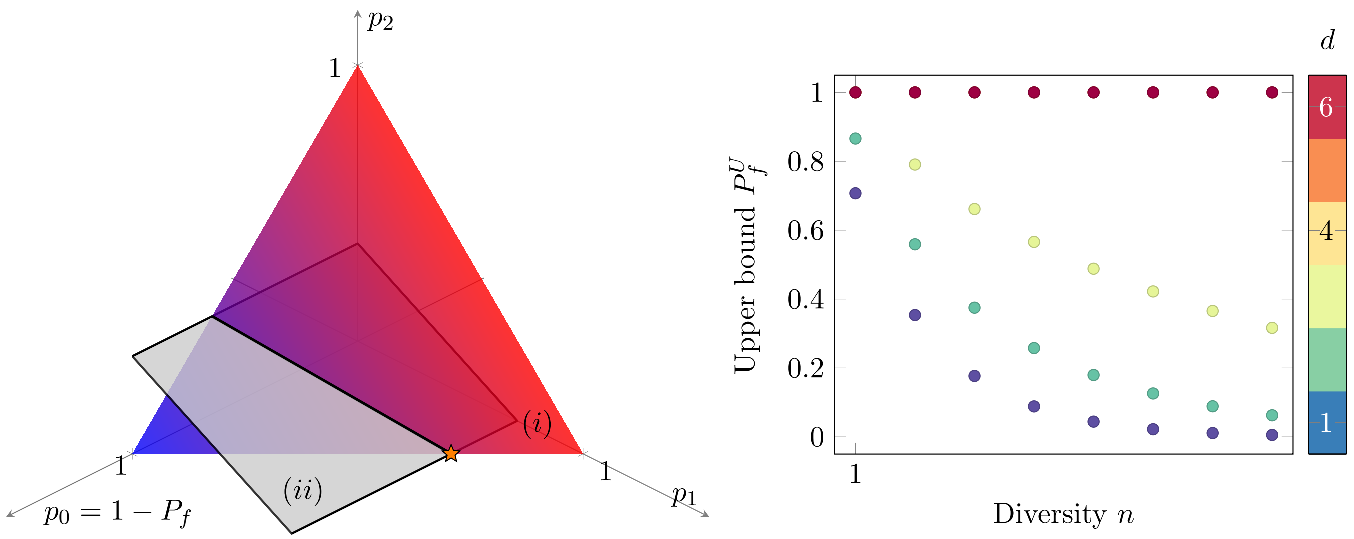


FIG. 6. **Using linear programming to obtain an exact upper bound of $P_f$.** (left) Geometric representation of optimizaion problem for $d = 2$, $n = 1$.Triangular plane corresponds to constraint (i) in Eq. 23. This plane is coloured according to the function we want to maximize, more red colours corresponding to higher values. The black plane parallel to the axis of $p_0$ corresponds to consraint (ii) in Eq. 23. The solution must lay in the intersection between the two planes, since only there both constraints are satisfied. Specifically, the optimal value is reached at the point marked with an orange star, since it is there where $P_f$ is maximum (most red part). (right) Exact uppeer bound of $P_f$ obtained through linear programming. Importantly $P_f$ cannot increase with $n$ when $d < 4$, because its upper bound decreases with $n$. On the contrary, for $d \geq 4$, the upper bound is always one, meaning that $P_f$ could increase with $n$.

(see Fig 4, left panel). That is, we maximize $P_f$ over the probability density vector $p$, subject to the constraints that $(i)$, the elements of $p$ sum to 1, $(ii)$, the expected value of the resulting probability distribution matches that in Eq. 12. To warm up, before considering the complete problem, we consider a simpler (only losely constrained) version of it. That is, we attempt to minimize the probability of feasibility over the variables $p_i$ (probability densities), only subjected to the constraint that the elements of the vector $p$ are nonnegative and add to 1.

$$\begin{aligned} \max_p \quad & 1 - p_0 \\ \text{s.t.} \quad & \sum_{i=0}^{d^n} p_i = 1 \quad . \end{aligned}$$

This problem has infinite solutions of the form

$$p^\star = (0, p_1, \dots, p_{d^n}) \quad . \tag{22}$$

That is, put the weight anywhere but on $p_0$, such that $P_f = 1$ (see Eq. 17). In Fig. 6**A**, this would correspond to solutions laying on the right edge of the triangular plane. Note that the maximizer of $1 - p_0$ is also the minimizer of $p_0$, which is the problem we solve.

Next, we introduce the constraint on the expected value of the distribution $P(N_+)$, which we know is given by Eq. 12. The minimization problem can now be stated as

$$\begin{aligned} \min_{p} \quad & p_0 \\ \text{s.t.} \quad & (i) \sum_{i=0}^{d^n} p_i = 1 \\ & (ii) \sum_{i=0}^{d^n} i p_i = \frac{\sqrt{d^n}}{2^n} \quad , \end{aligned} \tag{23}$$

where $\mathbb{E}[N_+]$ is given by Eq. 12.

The geometric representation of Eq. 23 is plotted in Fig. 6, left panel. We perform this minimization in general, for a range of $n$ and $d$ values, and plot the results in Fig. 6. We observe that $P_f^U$ is decreasing when $d < 4$, and equal to 1 when $d \geq 4$. This implies that $P_f$ has to be strictly decreasing in $n$ when the interaction order is less than 4, but could potentially increase otherwise.

The bound confirms that feasibility must fall with diversity when $d < 4$ and may rise otherwise. We close by placing these counts in a broader mathematical framework.

#### IX. INTEGRAL GEOMETRY

We recast the root counts in the language of integral geometry, where Howard's formula expresses the expected number of equilibria as a product of expected degrees.

The Kostlan-Shub-Smale-Edelman identity (10) is part of a more general mathematical framework, called *integral geometry*. We refer to the seminal paper by Kostlan and Edelman [5] that explains how integral geometry is used in the context of random polynomials.

Each random polynomial $f$ defines a *random projective hypersurface*

$$X(f) := \{x \in \mathbb{RP}^n \mid f(x) = 0\}.$$

We denote its *expected degree* by

$$\mathrm{edeg}(f) := \mathbb{E}\left[\#(X(f)\cap L)\right] \quad ,$$

where $L$ is a random line, that is independent of $f$, and defined by choosing $n-1$ independent linear equations with independent standard Gaussian coefficients.

Let us consider $n$ independent random real homogeneous polynomials $f_1,\dots,f_n$ in $n+1$ variables. We assume that the distribution of each $f_i$ is *orthogonally invariant* meaning that satisfies $f_i(x)\sim f_i(Ox)$ for every orthogonal matrix $O\in O(n+1)$. For instance, the KSS ensemble satisfies this property.

Howard's integral geometry formula [7], generalized by Bürgisser and Lerario [3, Theorem 3.19], states that under this assumption $X(f_1)\cap\cdots\cap X(f_n)$ consists of finitely many points almost surely, and that

$$\begin{aligned} E[N_{\mathbb{R}}] &= \mathbb{E}\left[\#\left(X(f_1)\cap\cdots\cap X(f_n)\right)\right] \\ &= \mathrm{edeg}(f_1)\cdots\mathrm{edeg}(f_n) \quad . \end{aligned} \tag{24}$$

Using the same argument as for (12) we get for the expected number of feasible equilibria:

$$\mathbb{E}[N_+] = \frac{\mathrm{edeg}(f_1)\cdots\mathrm{edeg}(f_n)}{2^n} \quad . \tag{25}$$

More generally, we consider $k<n$ homogeneous polynomials in $n+1$ variables. This is the case when we are interested in states of the Lotka-Volterra models (main text Eq. 1) where only a subset of variables is stable. [3, Theorem 7.2] yields that $X(f_1)\cap\cdots\cap X(f_k)$ almost surely is $(n-k)$-dimensional and that

$$\mathbb{E}\left[\mathrm{vol}_{n-k}(X(f_1)\cap\cdots\cap X(f_k))\right] = \mathrm{vol}_{n-k}(\mathbb{RP}^{n-k})\cdot\mathrm{edeg}(f_1)\cdots\mathrm{edeg}(f_k) \quad .$$

This viewpoint applies to any orthogonally invariant ensemble, a generality we exploit in the final section.

#### X. EXPECTED NUMBER OF ZEROS OF ORTHOGONALLY INVARIANT ENSEMBLES

We compute the expected degree for general orthogonally invariant ensembles, recovering the KSS count, allowing unequal degrees, and contrasting it with the real Fubini–Study ensemble.

Recall that we call a random ensemble of polynomials $f$ in $n+1$ variables orthogonally invariant, if $f(x) \sim f(Ox)$ for every orthogonal matrix $O \in O(n+1)$. The KSS ensemble is orthogonally invariant.

A KSS system $f$ of degree $d$ has

$$\mathrm{edeg}(f) = \sqrt{d} \quad .$$

We therefore recover (10) from (24):

$$E[N_{\mathbb{R}}] = \mathbb{E}\left[\#\left(X(f_1) \cap \cdots \cap X(f_n)\right)\right] = d^{n/2} \quad ,$$

where $f_1, \ldots, f_n \overset{\text{iid}}{\sim}$ KSS of degree $d$.

Interestingly, the formula also works when the KSS polynomials have differing degrees. Suppose that we have $f_i \sim$ KSS of degree $d_i$. Then,

$$E[N_{\mathbb{R}}] = (d_1 \cdots d_n)^{\frac{1}{2}} \quad .$$

This is the case of Lotka-Volterra models (main text Eq. 1) when the dynamics of population $i$ has interaction order $d_i$ with the other populations. Here, we have for the number of positive solutions (25)

$$\mathbb{E}[N_+] = \frac{(d_1 \cdots d_n)^{\frac{1}{2}}}{2^n} \quad . \tag{26}$$

As a function of $n$, when $d_1 \cdots d_n$ grows faster/slower than $4^n$, this function is increasing/decreasing. For instance, taking $d_1 = 2d$ and $d_2 = \cdots = d_n = d$, we have that $\mathbb{E}[N_+]$ increases precisely when $d > 2$.

More generally, for $f_i \sim$ KSS of degree $d_i$, we have

$$\mathbb{E}\left[\mathrm{vol}_{n-k}(X(f_1) \cap \cdots \cap X(f_k))\right] = \mathrm{vol}_{n-k}(\mathbb{RP}^{n-k}) \cdot (d_1 \cdots d_k)^{\frac{1}{2}} \quad .$$

Again, the volume of positive solutions inside this is given by dividing the right-hand side by $2^n$.

What about expected volumes of other ensembles? Fyodorov, Lerario and Lundberg [6] give a detailed answer. All orthogonally invariant random polynomials can be parameterized as follows (see [5]). Let $g_{j,\ell}$, where $j \in J_\ell$, be the standard basis of spherical harmonics of degree $\ell$ on the sphere $S^n$. The nonnegative weights $p_d(d), p_d(d-2), \ldots,$ parameterize all invariant random homogeneous polynomial in $x = (x_1, \ldots, x_{n+1})$ of degree $d$ via

$$f(x) = \sum_{d-\ell \text{ is even}} p_d(\ell) \sum_{j \in J_\ell} T_{j,\ell} \, \|x\|^{d-\ell} \, g_{j,\ell}(x/\|x\|) \quad ,$$

where $\|x\|^2 = x_1^2 + \cdots + x_{n+1}^2$ and the $T_{j,\ell}$ are i.i.d. standard Gaussian random variables.

In [6, Theorem 1 & Section 8, Example 3] it is shown that for every $0 < h \leq 1$ and any integrable function $\psi : \mathbb{R}_{>0} \to \mathbb{R}$ with subgaussian tails, if we define the random ensemble $f$ via

$$p_d(\ell) = \frac{1}{d^h}\,\psi\left(\frac{\ell}{d^h}\right) \quad , \tag{27}$$

then we have

$$c_1\, d^h \leq \mathrm{edeg}(f) \leq c_2\, d^h \quad ,$$

where $c_1, c_2 > 0$ are constants (independent of $d$, but not of $n$).

For instance, the KSS ensemble has $h = \frac{1}{2}$. Another interesting example is the *real Fubini-Study ensemble* (see [6, Section 8, Example 2]), which is defined by taking a random polynomial uniformly from the unit sphere in the $L^2$-norm in the space of all homogeneous polynomials in $n+1$ variables of degree $d$. This ensemble has $h = 1$. Therefore, by the formula above, if $f_1, \ldots, f_n$ are i.i.d. real Fubini-Study of degree $d$, we have

$$c_1' d^n \leq E[N_{\mathbb{R}}] \leq c_2' d^n \quad ,$$

where $c_1', c_2' > 0$ are independent of $d$. In this case,

$$c_1' \left(\frac{d}{2}\right)^n \leq E[N_+] \leq c_2' \left(\frac{d}{2}\right)^n \quad .$$

Compared to our analysis in Section II, where we considered asymptotics in $n$, we can't immediately use the last formula to study how $E[N_+]$ grows as a function of $n$, because we don't know how the constants $c_1', c_2'$ depend on $n$. Nevertheless, we still get the asymptotic behavior in $d$: while the number of real/positive zeros of the KSS ensemble grows as $d^{n/2}$, the number of real/positive zeros of the real Fubini-Study ensemble grows as $d^n$. In general, if we have a system of polynomials from an ensemble defined by (27), then the number grows as $d^{h \cdot n}$. This is a qualitative difference in how these numbers grow.

#### XI. OTHER RANDOM ENSEMBLES

Integral geometry exploits orthogonal invariance. There are other random polynomial ensembles, that are not orthogonally invariant, but we still know their expected value of real roots.

Most notably, Mathis [9] presents a formula for the following ensemble. Suppose the degree $d$ can be written as $d = \delta \cdot n$. Then, consider the following (non-homogeneous) ensemble:

$$f(x) = \sum_{0 \le a_1, \ldots, a_n \le \delta} \sqrt{\binom{\delta}{a_1} \cdots \binom{\delta}{a_n}}\, T_a\, x_1^{a_1} \cdots x_n^{a_n},$$

where the $T_a = T_{a_1,\ldots,a_n}$ are i.i.d. standard Gaussian random variables. This ensemble is not orthogonally invariant (since it doesn't use all monomials of degree at most $d$), but it's invariant under reflection of the variables; which is needed for our argument that $E[N_+] = \frac{1}{2^n} E[N_{\mathbb{R}}]$. Mathis proves in [9, Proposition 4.36] that if we take independent $f_1, \ldots, f_n$ with $\deg(f_i) = d_i$ from this ensemble one has

$$E[N_{\mathbb{R}}] = \frac{n!\, b_n}{(2m)^m} \sqrt{d_1 \cdots d_n},$$

where $b_n = \frac{\pi^{n/2}}{\Gamma(n/2+1)}$ is the volume of the standard Euclidean unit ball in $\mathbb{R}^n$.

#### XII. DISTRIBUTION OF THE JACOBIAN OF A KOSTLAN SYSTEM

Feasibility is necessary for coexistence but not sufficient; whether a feasible equilibrium is stable is decided by the spectrum of the community matrix, which at a root $x$ of the dynamics equals $\operatorname{diag}(x)\, Jf(x)$. Since the equilibria are themselves the random roots of $f$, the relevant object is the Jacobian at a fixed point $x$. Here we show this conditioned Jacobian is again Gaussian and compute its covariance, providing the starting point for a stability analysis of feasible high-$d$ equilibria.

Fix a point $x = (x_1, \ldots, x_n) \in \mathbb{R}^n$. Consider a system $f = (f_1, \ldots, f_n)$ of i.i.d. Kostlan polynomials. What is the distribution of

$$Jf(x) \mid \big(f(x) = 0\big),$$

i.e., what is the distribution of the Jacobian

$$Jf(x) = \left(\frac{\partial f_i}{\partial x_j}\right)_{1 \le i,j \le n}$$

conditioned on the event that $x$ is a zero of $f$. Since this event has probability zero, we need to make sense of it: $\{f \mid f(x) = 0\}$ is a linear space in the space of polynomial systems. A Gaussian random variable restricted to a linear space is still Gaussian. We are interested in the distribution of $Jf(x)$ with respect to the Gaussian distribution on $\{f \mid f(x) = 0\}$.

To compute this we first view $x$ as a point

$$[x_1 : \cdots : x_n : 1] \in \mathbb{RP}^n,$$

and the $f_i$ as homogeneous polynomials in $n+1$ variables. Let also

$$x' = \frac{(x_1, \ldots, x_n, 1)^T}{\sqrt{x_1^2 + \cdots x_n^2 + 1}}$$

be the corresponding point on the sphere. Writing

$$\tilde{J}f(x) = \left(\frac{\partial f_i}{\partial x_j}\right)_{1 \le i \le n, 1 \le j \le n+1}$$

for the projective Jacobian we have

$$Jf(x) = \tilde{J}f(x)\, P, \tag{28}$$

where

$$P = \begin{pmatrix} I_n \\ 0 \end{pmatrix} \in \mathbb{R}^{(n+1)\times n}$$

is the projection matrix that projects onto the first $n$ columns.

There is an orthogonal matrix $O \in O(n+1)$ with

$$Ox' = (0, \ldots, 0, 1) =: e.$$

Write $g(x) = f(O^T x)$. Since $f$ is Kostlan, also $g$ ist Kostlan. We have

$$\tilde{J}g(e) = \tilde{J}f(O^T e)\, O^T = \tilde{J}f(x')\, O^T. \tag{29}$$

Furthermore,

$$g(x) = x_{n+1}^d\, a + x_{n+1}^d \sqrt{d}\, A(x_1, \ldots, x_n)^T + \cdots\,,$$

where the dots on the right mean lower-order terms. Since $f(x) = 0$, we have $g(e) = 0$, which is equivalent to $a = 0$. The matrix $A \in \mathbb{R}^{n\times n}$ has i.i.d. standard normal entries, because $g$ is Kostlan. We obtain:

$$\tilde{J}g(e) = \begin{pmatrix} \sqrt{d}A & 0 \end{pmatrix} = \sqrt{d}A\, P^T.$$

By (29):

$$\tilde{J}f(x') = \tilde{J}g(e)\, O = \sqrt{d}A\, P^T\, O.$$

The law of linearly transforming Gaussians yields

$$Jf(x') \sim N(0, d\,\Sigma'),$$

where

$$\Sigma' = I_n \otimes (O^T P P^T O)$$

Since the entries of $\tilde{J}f(x')$ are homogeneous polynomials of degree $d-1$, setting $\lambda = \sqrt{x_1^2 + \cdots + x_n^2 + 1}$ we have

$$\tilde{J}f(x) = \tilde{J}f(\lambda\, x') = \lambda^{d-1} \tilde{J}f(x') \sim N(0, \lambda^{2(d-1)}\, d\,\Sigma').$$

Finally, by (28)

$$JF(x) = \tilde{J}F(x)\, P \sim N(0, \lambda^{2(d-1)}\, d\,\Sigma),$$

where

$$\begin{aligned}
\Sigma &= (I_n \otimes P^T)\Sigma'(I_n \otimes P) \\
&= I_n \otimes (P^T O^T P P^T O P) \\
&= I_n \otimes (U^T U), \qquad U = P^T O P.
\end{aligned}$$

Note that $\Sigma = \Sigma(x)$ depends on $x$, because $O$ depends on $x$. We have proved the following result.

**Theorem:** $Jf(x) \mid (f(x) = 0)$ is a random matrix that has centered Gaussian entries with covariance matrix given by $d\,\lambda^{2(d-1)}\,\Sigma(x)$.

#### APPENDIX

This appendix collects supporting numerical experiments that probe how robust the feasibility results are to relaxing the KSS assumptions.

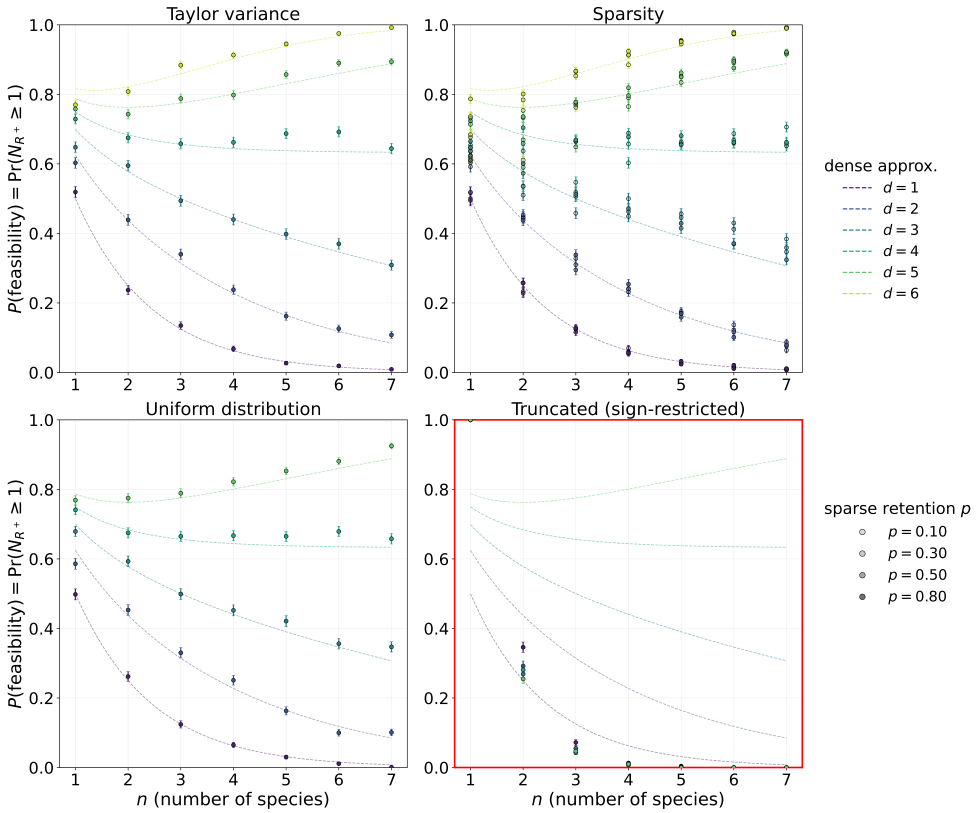


FIG. 7. Probability of feasibility $P_f = \Pr(N_+ \geq 1)$ as a function of population number $n$ for the four relaxations of the KSS ensemble: Taylor variance schedule (top-left), per-coefficient Bernoulli sparsity (top-right), Uniform coefficient distribution (bottom-left), and sign-restricted Truncated draws (bottom-right, framed in red). In each panel, color encodes the polynomial degree $d \in \{1, \ldots, 6\}$ and dashed lines overlay the KSS dense independent-root approximation $P_f \approx 1 - (1 - 2^{-n})^{d^{n/2}}$. The Taylor and Uniform panels track the KSS curve closely, confirming that swapping the variance schedule or the coefficient family (at fixed second moment) preserves the KSS feasibility scaling. The sparse panel encodes retention probability $p \in \{0.10, 0.30, 0.50, 0.80\}$ via marker opacity; deviations from the dense reference grow as $p$ decreases. The truncated panel is framed in red because sign-restriction breaks the KSS pattern: feasibility collapses to near zero for all but the smallest $(n, d)$. Markers are Monte Carlo estimates over $K = 1000$ trials per cell (error bars: $\pm 1\,\mathrm{SE}$).

[8] Lechón-Alonso, P., Z. R. Miller, A. Liaghat, P. Breiding, M. Pascual, and S. Allesina (2026), "Tipping points are typical in ecosystems with higher-order interactions," .

[9] Mathis, L. (2025), "Real gaussian exponential sums via a real moment map," arXiv:2411.11345 [math.PR].
